# Rad and Phospholamban are Key Drivers of the Ventricular Adrenergic Response and Stress-Induced Arrhythmia

**DOI:** 10.64898/2026.09.22.747614

**Authors:** Achal M. Gowda, Erick B. Ríos Pérez, Daniela Ponce-Balbuena, Samuel L. Shenkenberg, Francisco J. Alvarado

## Abstract

The adrenergic response is a fundamental mechanism that regulates heart rate (chronotropy), cardiac contractility (inotropy) and relaxation (lusitropy).^1^ Adrenergic stress is also a recognized trigger of arrhythmia in disease.^2^ Yet, our understanding of the underlying molecular basis remains incomplete. Protein kinase A (PKA) and the Ca^2+^/calmodulin-dependent kinase II (CaMKII) phosphorylate multiple targets proposed to participate in the adrenergic response, including the GTP-binding protein Rad, phospholamban (PLB) and ryanodine receptor 2 (RyR2).^1, 3–5^ Here we demonstrate that phosphorylation of both Rad and PLB is necessary for inotropy and lusitropy. We show that changes in cardiac contractility and relaxation are primarily dependent on intracellular Ca^2+^ handling. Finally, we report that Rad and PLB control stress-induced arrhythmogenesis, despite the phosphorylation of other pro-arrhythmic targets. We have identified the essential molecular components of the adrenergic response, resolving a long-standing debate in cardiac excitation-contraction coupling and refining current models of sympathetic regulation in health and disease.

## Main

Cardiac function is regulated by the sympathetic nervous system (SNS) through the adrenergic or “fight-or-flight” response. Epinephrine and norepinephrine, two catecholamines released during sympathetic stimulation, activate β_1_-adrenergic receptors (β_1_-AR) in the heart, leading to increased heart rate (chronotropy), contractility (inotropy) and rate of relaxation (lusitropy), among others.^1^ Chronotropy is regulated at the sinus node via specialized mechanisms of the coupled clock system, while inotropy and lusitropy are regulated in the ventricles through excitation-contraction coupling (ECC).^1, 6, 7^ In disease, adrenergic stress is a known trigger of life-threatening cardiac arrhythmias.^2^ Although this fundamental pathophysiological response has been studied extensively, our understanding of the underlying mechanisms remains incomplete, providing limited insights into how the system can fail in disease.

ECC is powered by intracellular Ca^2+^ cycling. Action potentials activate L-type Ca^2+^ channels (LTCCs), allowing a small influx of Ca^2+^ (*I*_CaL_) into the cardiomyocytes. *I*_CaL_ induces further Ca^2+^ release from the sarcoplasmic reticulum (SR), an intracellular store, via ryanodine receptor 2 (RyR2). The resulting increase in intracellular [Ca^2+^], or Ca^2+^ transient (CaT), activates the myofilaments and produces contraction. Relaxation results mainly from extrusion of Ca^2+^ from the cell through the Na^+^/Ca^2+^ exchanger 1 (NCX1) and reuptake into the SR by the sarco-(endo)-plasmic reticulum Ca^2+^ ATPase 2 (SERCA2) pump. Binding of catecholamines to β_1_-AR in ventricular myocytes increases cytosolic cAMP, activates protein kinase A (PKA) and ultimately enhances ECC. PKA phosphorylates three Ca^2+^ handling proteins (Fig. 1a): (1) the LTCC inhibitor Rad, increasing *I*_CaL_ density;^4^ (2) the SERCA2 inhibitor phospholamban (PLB), increasing SR Ca^2+^ uptake;^3, 5^ and (3) RyR2, which is proposed to modulate SR Ca^2+^ release.^8, 9^ The Ca^2+^/calmodulin-dependent kinase II (CaMKII) is also activated during adrenergic stimulation and phosphorylates many of the same molecular targets.^10^ Together, these changes lead to larger and faster CaTs that drive stronger and faster contractions. The hierarchy of these molecular contributors remains unsettled despite extensive research, and several reports have cast doubt on whether RyR2 phosphorylation contributes substantively to these dynamics.^9, 11–13^ In this study, we aimed to identify the key regulators of ventricular Ca^2+^ handling during adrenergic activation and to characterize how their individual and combined actions modulate cardiac function and arrhythmia susceptibility.

**Fig. 1.**
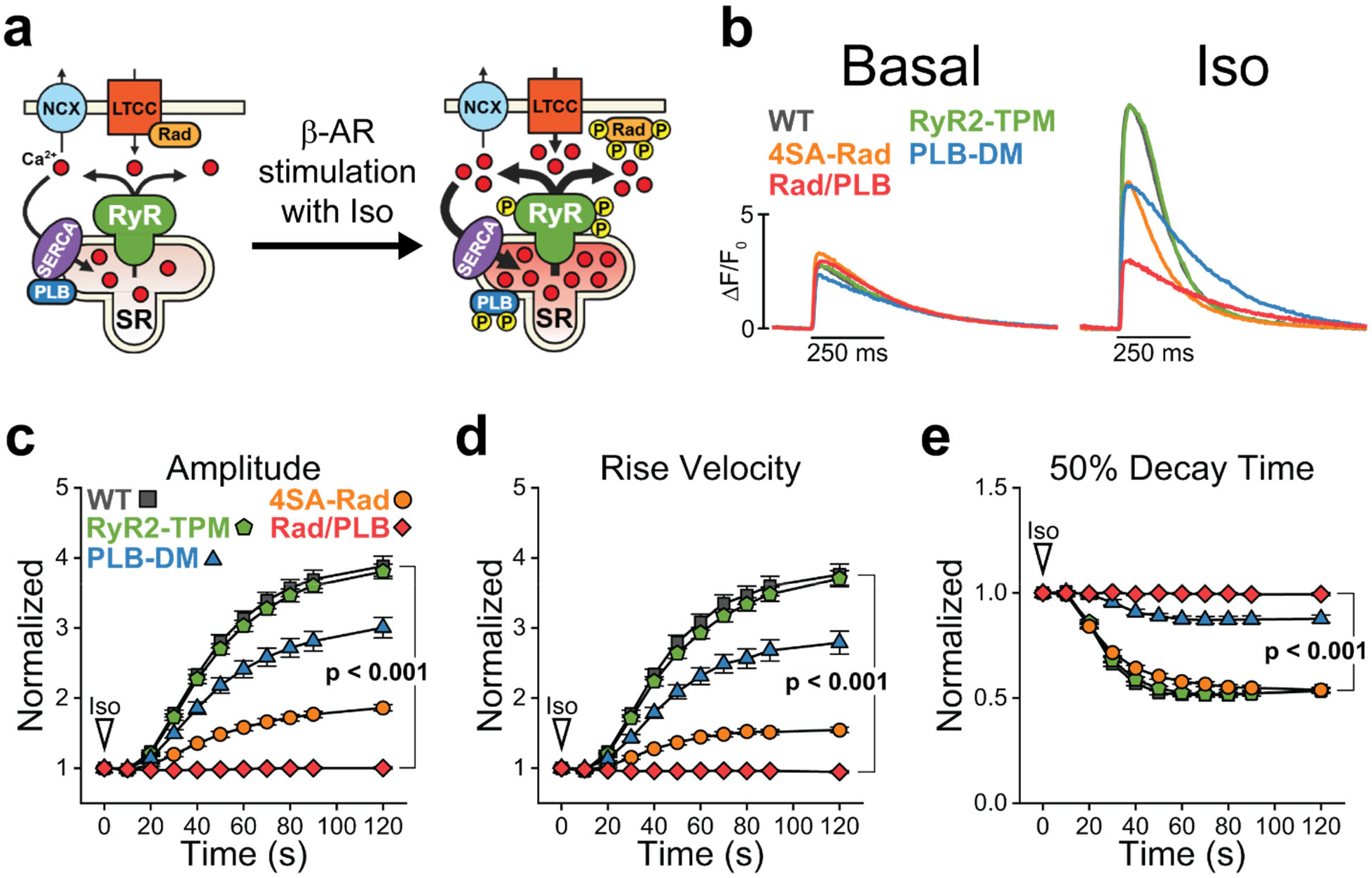
Phosphorylation of key proteins modulates the Ca^2+^ handling dynamics during β-adrenergic stimulation. **a.** Diagram of the major Ca^2+^ handling proteins regulated downstream of β-AR activation. GTP-binding protein Rad, phospholamban (PLB), and ryanodine receptor 2 (RyR2) are phosphorylated, leading to increased *I*_CaL_, increased SR Ca^2+^ release, faster Ca^2+^ release and reuptake, and increased SR Ca^2+^ load. **b.** Representative confocal CaT fluorescence profiles obtained in ventricular cardiomyocytes at 1 Hz pacing after 120 s under basal conditions or in the presence of 100 nM isoproterenol (Iso). Data are presented as corrected fluorescence (ΔF/F_0_). **c-e.** Time-course of CaT properties during Iso stimulation, normalized to basal values for each group: CaT amplitude (**c**), maximum rise velocity (**d**), and 50% decay time (**e**). Myocytes paced at 1 Hz for 30 seconds in basal conditions, followed by 120 seconds under 100 nM Iso perfusion. N = 6 mice per genotype. n = 56 WT and Rad/PLB, 42 RyR2-TPM, 30 4SA-Rad and PLB-DM myocytes. Linear mixed-effects model (Bonferroni test). P < 0.001 between WT and Rad/PLB for all parameters.

We began by evaluating Ca^2+^ handling in cardiomyocytes from mice carrying phospho-ablations at key residues targeted during the adrenergic response: RyR2-S2031A/S2808A/S2814A (triple phospho-mutant, RyR2-TPM),^11^ Rad-S25A/S38A/S272A/S300A (4SA-Rad)^4^ and PLB-S16A/T17A (PLB-DM).^5^ CaTs were elicited with field stimulation in basal conditions and under perfusion with 100 nM isoproterenol (Iso), a β-adrenergic agonist, to assess global Ca^2+^ handling under confocal imaging. As we previously reported, RyR2-TPM myocytes, which lack the known PKA and CaMKII sites, showed CaT dynamics nearly indistinguishable from WT controls in basal conditions or under adrenergic stimulation (Fig. 1b-e, Extended Data Fig. 1). While phosphorylation of RyR2 has been proposed to be critical to achieving a complete adrenergic response,^8, 9, 14–16^ numerous studies have contested these results.^13, 17–21^ Our data reinforce the idea that phosphorylation of RyR2 at these “canonical” sites is dispensable for global Ca^2+^ handling during adrenergic stimulation.^11^ In contrast, targeting the phospho-sites in Rad and PLB significantly blunted different properties of the adrenergic response. 4SA-Rad myocytes, which lack PKA-mediated regulation of *I*_CaL_,^4^ had a reduced CaT response to Iso stimulation compared to WT controls, including CaT amplitude, rise velocity and SR content (Fig. 1b-e, Extended Data Fig. 2). However, CaT decay and rate of SR Ca^2+^ reuptake were unaffected (Fig. 1e, Extended Data Fig. 2e,g). PLB-DM myocytes, which lack PKA-and CaMKII-mediated regulation of SR Ca^2+^ uptake via SERCA2, also showed a lower Ca^2+^ handling response to adrenergic stimulation, with reduced CaT amplitude, rise velocity and SR load than controls (Fig. 1b-e, Extended Data Fig. 3).^3, 5^ PLB-DM cells also showed lower basal SR load and diminished response in CaT decay and SR Ca^2+^ reuptake, consistent with persistent inhibition of SERCA2 by PLB (Fig. 1e, Extended Data Fig. 3e-g).

Given that both 4SA-Rad and PLB-DM myocytes showed a residual adrenergic response, notably in CaT amplitude and velocity and SR load, we crossbred both strains to create a model harboring all six phospho-ablations (4SA-Rad/PLB-DM or Rad/PLB, Fig. 2a). Remarkably, Iso failed to increase CaT amplitude and accelerate CaT rise and decay in Rad/PLB myocytes (Fig. 1c-e, Fig. 2b-f). An assessment of Ca^2+^ sinks and sources revealed that Rad/PLB myocytes cannot increase the cellular Ca^2+^ influx and retention that characterize the adrenergic response. Basal SR load was lower in Rad/PLB myocytes compared to WT controls, and SR Ca^2+^ content remained unchanged during Iso perfusion (Fig. 2g). The rate of SR Ca^2+^ uptake increased in WT cardiomyocytes treated with Iso, as expected, while Rad/PLB myocytes displayed a slight, yet significant decrease (Fig. 2h). Rad/PLB cardiomyocytes remained unresponsive to β-adrenergic stimulation at Iso concentrations ranging from 50 nM to 1 µM (Extended Data Fig. 4a-e). Interestingly, non-SR cytosolic Ca^2+^ removal (via NCX1, the plasma membrane Ca^2+^ ATPase [PMCA] and capture by other organelles such as the mitochondria^22^) increased significantly in the Rad/PLB group under Iso in a dose-dependent manner, while remaining unchanged in WT controls (Fig. 2i, Extended Data Fig. 4). Ca^2+^ extrusion via NCX1 is not believed to respond to β-adrenergic stimulation by Iso, and the contributions of PMCA and the mitochondria to CaT decay are thought to be limited.^22, 23^ Hence, the increase in non-SR-mediated Ca^2+^ removal with Iso in Rad/PLB cardiomyocytes suggests the presence of an adrenergically regulated Ca^2+^ removal pathway that will warrant further examination. Finally, *I*_CaL_ remained unchanged under Iso stimulation in Rad/PLB myocytes (Extended Data Fig. 5). These data indicate phosphorylation of both Rad and PLB is required for the Ca^2+^-handling component of the ventricular adrenergic response.

**Fig. 2.**
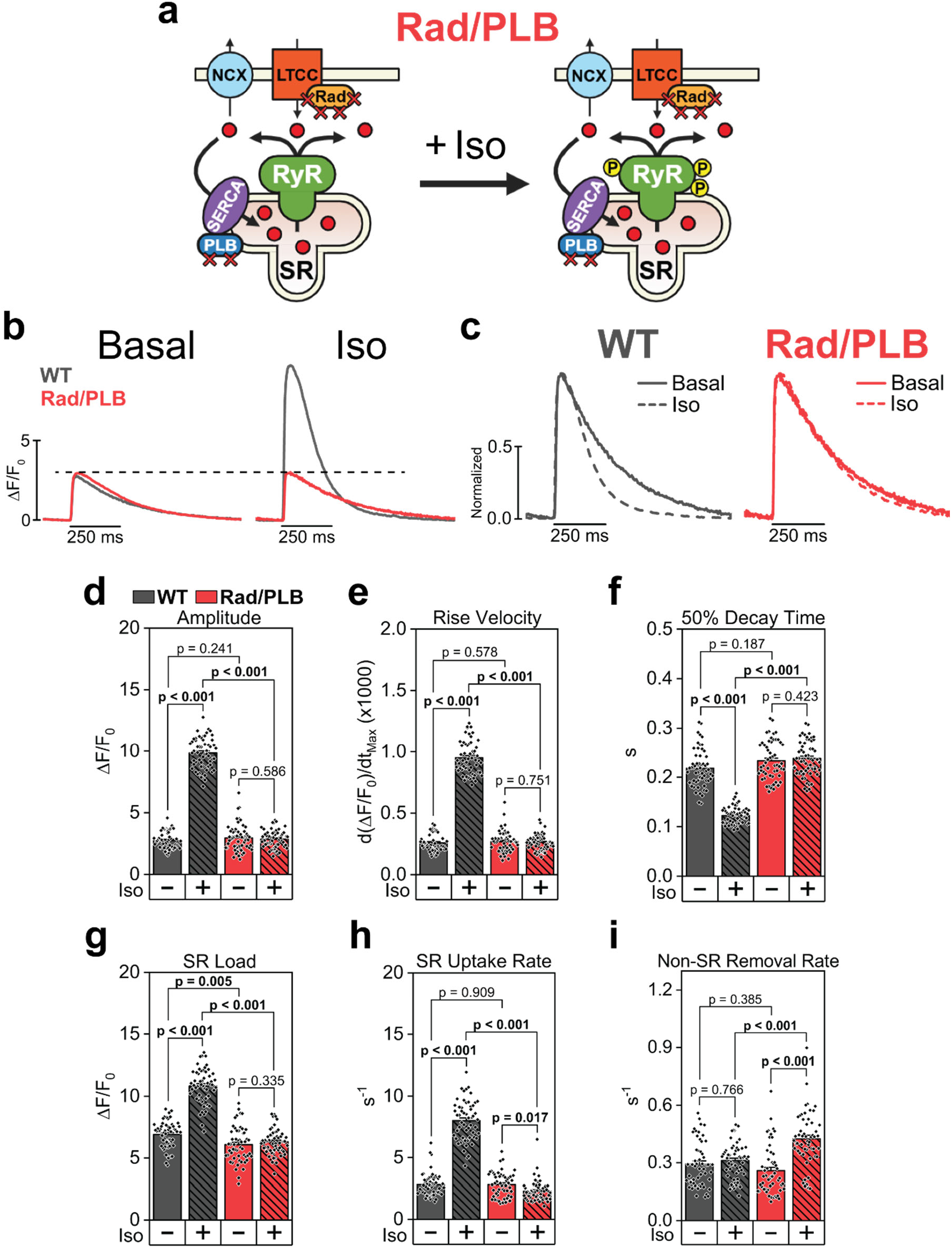
β-adrenergic control of Ca^2+^ dynamics is abolished in cardiomyocytes lacking critical phosphorylation sites in Rad and PLB. **a.** Diagram of Ca^2+^ handling response to β-AR activation by Iso in Rad/PLB cardiomyocytes. Phosphorylation sites on Rad and PLB are ablated, preventing normal adrenergic modulation of CaT properties. **b-c.** Representative confocal CaT fluorescence profiles obtained in wild-type (WT) and Rad/PLB ventricular cardiomyocytes at 1 Hz pacing after 120 s under basal conditions or in the presence of 100 nM Iso. Data are presented as corrected fluorescence (ΔF/F_0_) (**b**) or normalized to the peak of the CaT for each group (**c**). **d-i.** Quantification of CaT properties at 1 Hz pacing, after 120 s in basal conditions or under 100 nM Iso perfusion, including CaT amplitude (**d**), maximum rise velocity (**e**), 50% decay time (**f**), measured as the time from the CaT peak to 50% CaT decay, SR load (**g**), measured as the peak of a caffeine-induced CaT at the end of the CaT recording protocol, SR Ca^2+^ uptake rate (**h**), and non-SR Ca^2+^ removal rate (**i**). A single exponential fitting of the decay phase of the caffeine-induced transient was used to determine non-SR Ca^2+^ removal, while the SR Ca^2+^ uptake rate was calculated as the constant from exponential fitting of paced CaT decay minus the non-SR removal constant. N = 6 mice per genotype for all panels. n = 56 WT and Rad/PLB [Basal and Iso] myocytes for panels d-f. n = 55 WT [Basal], 54 WT [Iso], 54 Rad/PLB [Basal], and 55 Rad/PLB [Iso] myocytes for panel g. n = 54 WT and Rad/PLB [Basal and Iso] myocytes for panels h-i. Linear mixed-effects model (Bonferroni test) used for all comparisons.

Despite this major disruption to the cardiac adrenergic response, Rad/PLB hearts do not display an overt basal phenotype. Cardiac expression of essential Ca^2+^-handling proteins including RyR2, the pore-forming subunit of the LTCC Ca_V_1.2, NCX1, and SERCA2, was comparable to that of WT controls (Extended Data Fig. 6a-b). PLB expression was increased in Rad/PLB hearts, which was also observed in the PLB-DM model.^5^ Given that basal CaT decay and SR uptake rate were unaffected in Rad/PLB myocytes (Fig. 2f,h), it is unlikely that higher PLB expression directly affects Ca^2+^ handling or cardiac function. However, the enhanced inhibitory effect of PLB on SERCA2 became evident in Rad/PLB myocytes under Iso, as demonstrated by the significantly reduced SR uptake rate (Fig. 2h). At the whole-animal level, echocardiography showed normal heart structure in Rad/PLB mice (Extended Data Table 1), although they displayed a tendency towards left ventricular (LV) dilation, similar to the 4SA-Rad mouse line.^4^ Nevertheless, Rad/PLB mice have normal cardiac contractility in basal conditions, with LV ejection fraction, fractional shortening, and stroke volume similar to that of WT mice (Extended Data Table 2).

During β-adrenergic stimulation, PKA phosphorylates serines 2031 and 2808 on RyR2, while CaMKII phosphorylates mainly serine 2814 and, to a lower extent, serine 2808 (Extended Data Fig. 6c).^8, 9, 15, 24^ Iso prompted phosphorylation of the three residues in both WT and Rad/PLB hearts (Extended Data Fig. 6d-g). However, we observed a significantly higher level of phosphorylation at S2031 under Iso in Rad/PLB hearts compared to WT controls (Extended Data Fig. 6d-e). In contrast, phosphorylation of S2814 was slightly reduced in Iso-treated Rad/PLB hearts (Extended Data Fig. 6d,g). As CaMKII is primarily a Ca^2+^-dependent kinase, we surmise that enzyme activation is limited by the lack of intracellular Ca^2+^ augmentation in Rad/PLB cardiomyocytes. Interestingly, both PKA and CaMKII can phosphorylate RyR2 at residues other than the three canonical sites (Extended Data Fig. 7), as has been noted in previous studies.^25^ This suggests there may be uncharacterized phosphorylation sites that could contribute to RyR2 regulation during the adrenergic response. Critically, we took an agnostic approach by using the Rad/PLB model: with intact RyR2 channels, any possible contribution of the channel to the global adrenergic response can be evaluated independently of specific residues. Hence, RyR2 phosphorylation does not appear to have a major contribution to the regulation of global Ca^2+^ cycling under adrenergic stimulation.

We next measured sarcomere contraction and relaxation kinetics in isolated ventricular cardiomyocytes. WT cells showed the expected increase in sarcomere contractility and contraction/relaxation velocity, but these parameters remained unchanged after Iso perfusion in Rad/PLB cells (Fig. 3a-d). Hence, the cellular inotropic and lusitropic responses were completely abolished in Rad/PLB myocytes. Alongside the Ca^2+^ handling apparatus, PKA and CaMKII phosphorylate myofilament proteins which are also proposed to influence adrenergic inotropy and lusitropy. Phosphorylation of troponin I decreases myofilament affinity for Ca^2+^, which may contribute to positive lusitropy by accelerating Ca^2+^ unbinding.^26^ Conversely, MyBP-C phosphorylation may impact inotropy through regulation of cross-bridge cycling.^27^ Given that we did not observe a positive response in Rad/PLB myocytes, it is possible that the contributions of myofilament protein phosphorylation are secondary to the enhancement of Ca^2+^ cycling. We also assessed the adrenergic response in anesthetized mice with echocardiography, as done previously.^11^ Iso elicited a typical positive inotropic response in WT hearts, with significant increases in LV ejection fraction (EF) and fractional shortening (FS) (Fig. 3e-g). Surprisingly, Rad/PLB mice showed a significant decrease in both parameters, along with a decrease in stroke volume (Extended Data Table 2). This may indicate extra-cardiac effects of Iso, such as β_2_-AR-mediated vasodilation, uncovered in the absence of positive inotropy and lusitropy.

**Fig. 3.**
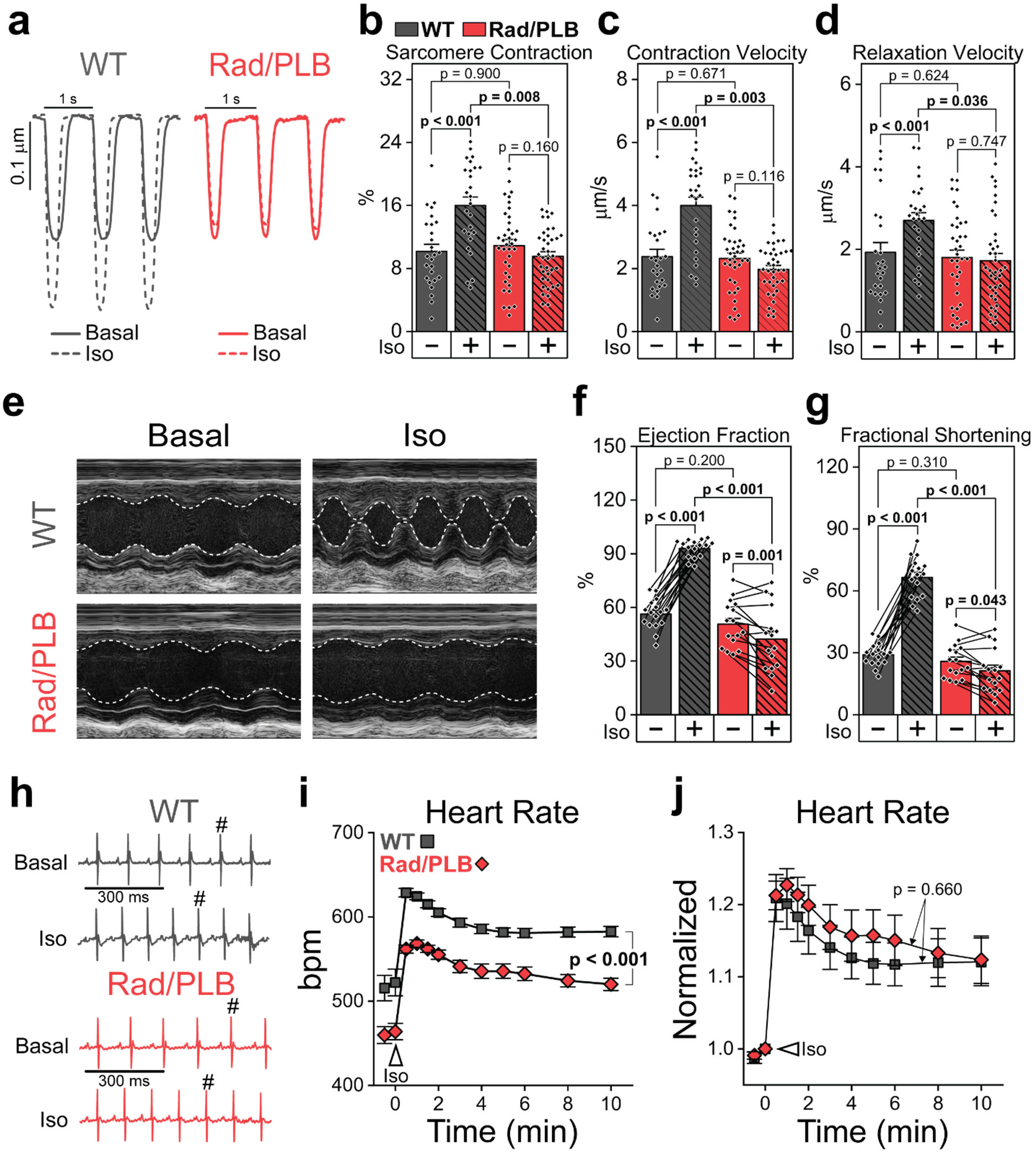
Positive inotropy and lusitropy are eliminated in the Rad/PLB model, while positive chronotropy is preserved. **a.** Representative traces of sarcomere contraction and relaxation in isolated cardiomyocytes, elicited with 1 Hz field stimulation and recorded before and after application of Iso (100 nM). **b-d**. Quantification of sarcomere shortening as a percentage of diastolic length (**b**), maximum contraction velocity (**c**), and maximum relaxation velocity (**d**). N = 4 WT and 5 Rad/PLB mice. N = 26 WT and 37 Rad/PLB myocytes. Linear mixed-effects model (Bonferroni test) used for all comparisons. **e.** Representative M-mode echocardiography line-scan images of the left ventricle, taken before and 2 min. after i.p. injection of Iso (2 mg/kg). **f-g.** Quantification of left ventricular ejection fraction (**f**) and fractional shortening (**g**), before and 2 min. after Iso injection. N = 16 mice per genotype. Linear mixed-effects model (Bonferroni test) with repeated measures used for both comparisons. **h.** Representative surface electrocardiography (ECG) traces before and 2 min after i.p. injection of Iso (2 mg/kg). ^#^ 5^th^ beat of ECG trace marked for visualization of chronotropic effect. **i-j.** Time-course of the chronotropic response to Iso injection via quantification of heart rate, shown as absolute heart rate (**i**) and normalized to heart rate at time zero (**j**). N = 6 WT and 5 Rad/PLB mice. Linear mixed-effects model (Bonferroni test) with repeated measures used for both comparisons. P < 0.001 between WT and Rad/PLB curves for i, and p = 0.660 between WT and Rad/PLB curves for j.

We continued with an evaluation of the chronotropic response in anesthetized mice. Adrenergic acceleration of heart rate (HR) requires either Rad-mediated regulation of Ca_V_1.3, an LTCC isoform in the sinus node, or direct cAMP regulation of nucleotide-gated HCN4 channels.^28^ Hence, we hypothesized that targeting Rad and PLB would not interfere with cardiac chronotropy. Rad/PLB mice displayed significant basal bradycardia (Fig. 3h-i); however, they showed a significant positive chronotropy in response to Iso. At all times post-Iso injection, Rad/PLB mice had lower HR than WT mice (Fig. 3i), but the normalized response was comparable between genotypes (Fig. 3j). Preventing Rad phosphorylation will interfere with adrenergic regulation of Ca_V_1.3,^28^ and blocking PLB phosphorylation will affect the Ca^2+^ clock.^29^ Both mechanisms appear to participate in determining automaticity in the sinus node and HR. However, intact cAMP regulation of HCN4 in Rad/PLB hearts can drive a positive chronotropic response of equal magnitude to WT controls.^28^

Surges of adrenergic stimulation during stress can induce cardiac arrhythmia and sudden death in different forms of heart disease. A prominent cellular trigger involves spontaneous SR Ca^2+^ release (SCR) through hyperactive RyR2 channels. Mutations and post-translational modifications in RyR2 contribute to channel hyperactivity, while Ca^2+^ loading brings SR content to threshold levels that promotes SCR.^30, 31^ However, it remains unresolved whether both conditions are necessary for arrhythmogenesis. Because Rad/PLB mice lack adrenergic regulation of cardiac Ca^2+^ handling, we hypothesized they would be protected from stress-induced arrhythmias. To test this, we employed a pharmacological challenge with epinephrine and caffeine commonly used in anesthetized mice.^11, 32^ We used a high dose (4 mg/kg Epi + 120 mg/kg caffeine), which consistently induced ventricular tachycardia (VT) in WT mice (Fig. 4). Remarkably, Rad/PLB mice showed a stable electrocardiogram without signs of sustained ventricular ectopy (Fig. 4), remaining resistant to VT during the challenge. These data suggest that Rad and PLB are critical for stress-induced arrhythmogenesis. Other pro-arrhythmic substrates such as RyR2 are intact in Rad/PLB mice. Hence, sub-threshold SR loading under stress is the most likely mechanism suppressing arrhythmia in these animals. These observations may have broad implications in disease. For example, gain-of-function mutations in RyR2 lead to catecholaminergic polymorphic ventricular tachycardia (CPVT), a syndrome that predisposes patients to life-threatening arrhythmia during stress. Our results support the notion that constraining SR loading can suppress arrhythmias in CPVT even if the primary defect in RyR2 remains,^33^ underscoring Rad and PLB as potential therapeutic targets for CPVT. Validation of this paradigm will require further experimentation.

**Fig. 4.**
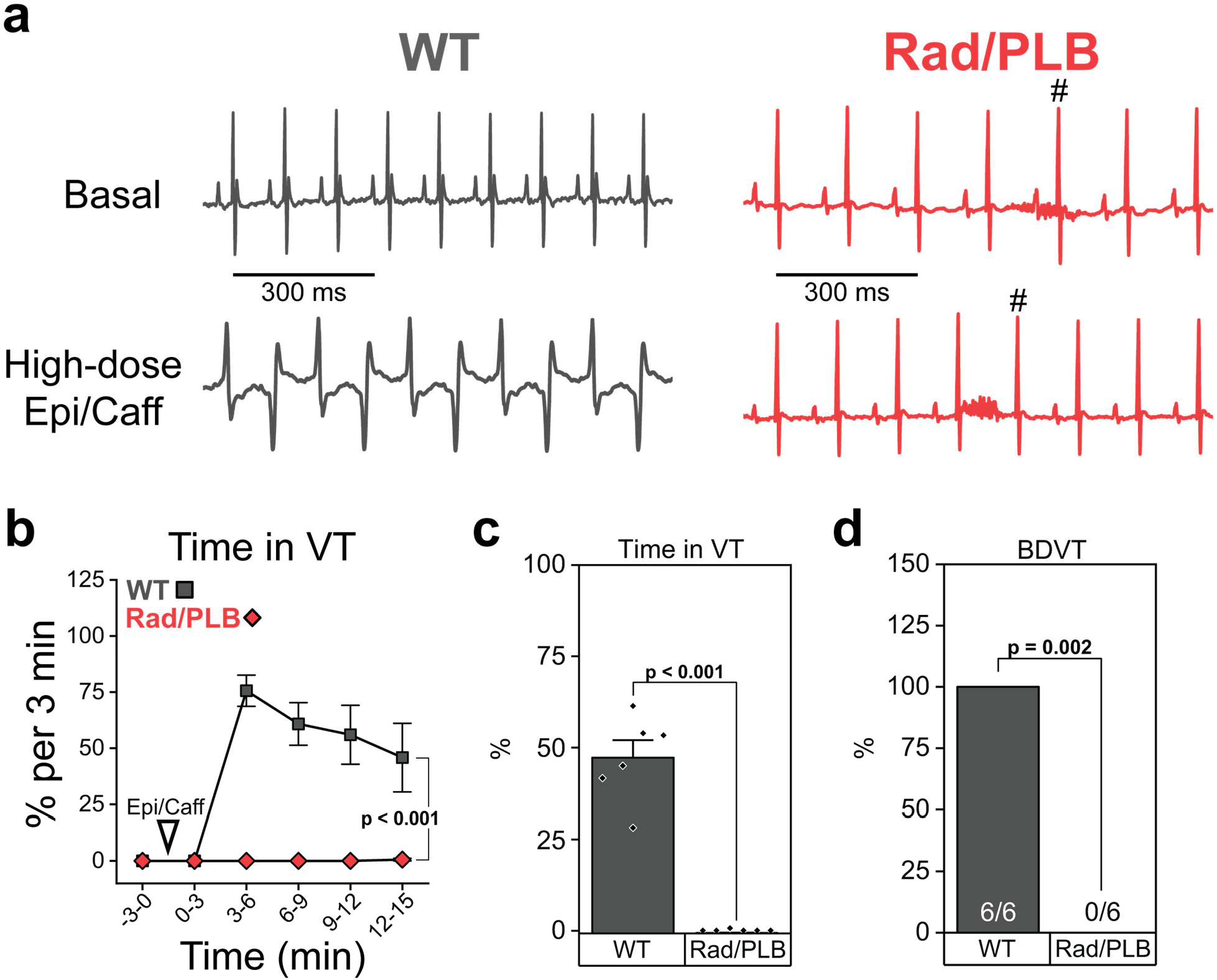
Rad and PLB are critical mediators of stress-induced arrhythmias. **a.** Representative surface electrocardiogram (ECG) traces before and after i.p. injection of epinephrine and caffeine (4 mg/kg and 120 mg/kg, respectively). WT mice consistently exhibit bidirectional polymorphic ventricular tachycardia (BDVT). Rad/PLB mice are resistant to ventricular tachyarrhythmias (VT) despite adrenergic activation (^#^5^th^ beat of ECG trace marked for visualization of chronotropic effect). **b.** Time course showing percent of time spent in VT in three-minute periods during the epinephrine/caffeine (Epi/Caff) challenge. Epi/Caff were injected at timepoint 0. N = 6 mice per genotype. Linear mixed-effects model (Bonferroni test) with repeated measures used for both comparisons. **c.** Total percent of time in VT throughout the 15-minute period post-injection of the Epi/Caff cocktail. N = 6 mice per genotype. T-test used for this comparison. **d.** Percent of mice exhibiting bidirectional ventricular tachycardia (BDVT) post-injection. N = 6 mice per genotype. Fisher’s exact test.

The molecular players involved in cardiac adrenergic regulation have complex overlapping functions, which has hindered the ability to assess the relative contributions of individual proteins. Here, we employ several knock-in mouse models with phospho-site ablations to dissect and characterize the molecular mechanisms of the ventricular adrenergic response. Using a model in which the three canonical phosphorylation sites on RyR2 are removed (RyR2-TPM), we reaffirm previous results from our laboratory suggesting that proposed critical RyR2 phospho-sites are dispensable in the CaT modulation during adrenergic stimulation (Fig. 1b-e, Extended Data Fig. 1).^11^ With phospho-null models of PLB (PLB-DM) and Rad (4SA-Rad), we confirm the contributions of SERCA2 regulation by PLB and LTCC regulation by Rad to β-adrenergic effects on Ca^2+^ dynamics (Fig. 1b-e, Extended Data Fig. 2, Extended Data Fig. 3).^4, 5^ To complete the systematic breakdown of the ventricular adrenergic response, the double-mutant Rad/PLB shows a remarkable elimination of Ca^2+^ handling changes in response to β-adrenergic stimulation (Fig. 1b-e, Fig. 2, Extended Data Fig. 4, Extended Data Fig. 5). Hence, we conclude that both Rad and PLB are the dominant regulators of inotropy and lusitropy, and each mechanism, on its own, can only trigger a partial response. This paradigm contrasts with that of chronotropy, in which either Rad/Ca_V_1.3 or cAMP/HCN4 can drive full heart rate acceleration independently of each other.^28^

Attempts to study the role of RyR2 phosphorylation in the adrenergic response have varied widely, from phospho-null and phospho-mimetic mutant models to voltage-clamped cardiomyocytes with controlled SR load.^8, 9, 14, 15, 34–37^ The conclusions of these studies were equally varied, resulting in vigorous debates over the importance of specific RyR2 phosphorylation sites, accessory proteins, etc. One of several complications with phospho-null models, which are the most widely used models, is that preventing phosphorylation of one residue can result in altered phosphorylation levels of remaining sites.^9, 17^ Because Rad/PLB myocytes are unresponsive to Iso and RyR2 phosphorylation is largely preserved, it is reasonable to conclude that RyR2 phosphorylation is dispensable for inotropy and lusitropy, settling a long-standing debate. Therefore, systolic SR Ca^2+^ release during adrenergic stimulation is graded by the combined effect of Rad and PLB, while RyR2 channels respond to the trigger *I*_CaL_ and release commensurately with SR content. This, however, does not preclude RyR2 from having a role in modulating diastolic sub-cellular Ca^2+^ dynamics, as we have proposed.^11^

By establishing Rad and PLB as the principal determinants of cardiac inotropy and lusitropy, our findings clarify the molecular architecture underlying two of the most critical physiological responses of the heart. Targeting a single protein in 4SA-Rad or PLB-DM mice blunts the response, but both models retain residual regulation of all properties of CaT dynamics and, therefore, of inotropy and lusitropy. The relative contribution of each protein can be further dissected based on our results: PLB regulation of SERCA2 is the dominant regulator of cytosolic Ca^2+^ removal and lusitropy, while Rad regulation of LTCC has a larger contribution to CaT amplitude and inotropy. Beyond normal physiology, adrenergic dysregulation is involved in a wide spectrum of cardiac pathologies, including arrhythmias, hypertrophy, and heart failure.^38–40^ We demonstrate that preventing adrenergic regulation of Ca^2+^ handling has an anti-arrhythmic effect, positioning Rad and PLB as potential pharmacological targets. By resolving key mechanisms of adrenergic control, this work provides a mechanistic foundation that may inform the development of more effective and precisely targeted therapies.

## Methods

### Study approval

Animal use was approved by the University of Wisconsin-Madison School of Medicine and Public Health Institutional Animal Care and Use Committee (M5944), which runs an animal program accredited by the Association for Assessment and Accreditation of Laboratory Animal Care International. Animal husbandry was performed by specialized personnel of Biomedical Research Model Services following standard procedures. Animals were housed in rooms with temperature and humidity control and under a 12-hour light/dark cycle. Veterinarians of the Research Animal Program and Compliance provided specialized care.

### Chemicals and reagents

All chemicals and reagents were obtained from MilliporeSigma unless noted otherwise.

### Generation of mouse models

RyR2-S2031A/S2808A/S2814A (RyR2-TPM) mice were previously generated in an 129S1/SvImJ background.^11^ 129S1/SvImJ RyR2-TPM mice were cross-bred with C57Bl/6J mice for >7 generations until the genetic background of the mice reached >99% C57Bl/6J. Rad-S25A/S38A/S272A/S300A (4SA-Rad) mice were obtained from the lab of Dr. Steven Marx at Columbia University.^4^ PLB-S16A/T17A (PLB-DM) mice were previously created using CRISPR-Cas9 technology by the University of Wisconsin-Madison Animal Models Core (RRID: SCR_024797) and Advanced Genome Editing Laboratory (RRID:SCR_021070).^5^ To generate 4SA-Rad/PLB-DM (Rad/PLB) mice, 4SA-Rad and PLB-DM mice were cross-bred until mice homozygous for all mutations were obtained. Mouse colonies were maintained by breeding homozygotes. Mice 12–24 weeks of age of both sexes were used for experiments. Age-and sex-matched C57Bl/6J mice were used as WT controls.

### Isolation of cardiomyocytes

Mice were injected with heparin (0.5 U g^−1^, intraperitoneal) and euthanized by cervical dislocation after 10 minutes. Hearts were quickly excised and the aorta was cannulated. Hearts were then mounted on a Langendorff apparatus and perfused with perfusion buffer (113 mM NaCl, 4.7 mM KCl, 1.2 mM MgSO_4_-7 H_2_O, 10 mM HEPES, 0.6 mM Na_2_HPO_4_, 12 mM NaHCO_3_, 0.6 mM KH_2_PO_4_, 10 mM KHCO_3_, 30 mM taurine, 500 mM 2,3-butanedione monoxime and 5.5 mM glucose with pH 7.40) at 37°C at approximately 3 ml min^−1^. Following several minutes of perfusion with normal perfusion buffer, hearts were perfused with perfusion buffer supplemented with 773.48 U ml^−1^ Collagenase Type II (Worthington) and 12.5 μM CaCl_2_. Once fully digested (5–7 minutes), the ventricles were excised and minced in perfusion buffer containing 10% FBS and 12.5 μM CaCl_2_. Tissue pieces were gently shaken and pipetted to dissociate cells, followed by filtration with a sterile mesh filter. Ca^2+^ was slowly reintroduced to dissociated cardiomyocytes in five steps for a final concentration of 1.8 mM Ca^2+^. Following Ca^2+^ reintroduction, cells were kept in normal Tyrode’s solution (135 mM NaCl, 4 mM KCl, 1.8 mM CaCl_2_, 1 mM MgCl_2_, 10 mM HEPES, 1.2 mM NaH_2_PO_4_, and 10 mM glucose, with pH 7.40) until used.

### Calcium imaging and analysis of Ca^2+^ transients

Confocal imaging of calcium was performed in isolated cardiomyocytes with a Zeiss LSM 980 confocal microscope. Myocytes were incubated with fluo-4 AM (10 μM dissolved in 0.4% Pluronic F-127, Thermofisher) for 5 minutes at 37 °C and then plated on laminin-coated 35 mm glass-bottom dishes (MatTek) in normal Tyrode’s solution. Ca^2+^ transients (CaTs) were elicited by field stimulation at 1 Hz (2 ms pulses, 60-100 V) and were recorded as follows: 1) 5 s without stimulation to record basal fluorescence (F_0_), 2) 30 s of 1 Hz stimulation with perfusion of normal Tyrode’s solution, 3) 120 s of 1 Hz stimulation with perfusion of either normal Tyrode’s solution or Tyrode’s solution with 100 nM Iso, 4) and a 5 s pulse of caffeine (10 mM, dissolved in Tyrode’s solution) to measure SR Ca^2+^ load. Fluorescence was excited at 488 nm and recorded at higher than 505 nm. Longitudinal line scans were recorded using a 40x/1.2 water immersion objective at 2.46 ms per line with ZEN 3.12 software (Carl Zeiss). Three Ca^2+^ transients were averaged at each time point over the 120 s period. All recordings were performed at room temperature. Transients were analyzed using custom MATLAB 2026b (MathWorks) scripts, as described before.^11^ SR Ca^2+^ uptake and non-SR Ca^2+^ removal rates were calculated from exponential fitting of the decay phase of the paced and caffeine-induced CaTs. While caffeine is present, persistently activating RyR2, SR Ca^2+^ sequestration is negligible. Therefore, the decay constant of the caffeine-induced transient (K_Caff_) represents the rate of Ca^2+^ removal by non-SR mechanisms (K_Caff_ = K_Non-SR_). During a normal paced CaT decay, Ca^2+^ removal is mediated by SERCA2 (SR Ca^2+^ uptake) and non-SR Ca^2+^ removal mechanisms (non-SR Ca^2+^ extrusion [NCX1, PMCA] and uptake [mitochondria and other organelles]). Therefore, the SR uptake rate can be estimated by subtracting K_Non-SR_ from the decay constant of the paced CaT (K_SR_ = K_CaT_ – K_Non-SR_).^41^

### Homogenization of cardiac tissue and western blotting

WT and Rad/PLB mice were euthanized by cervical dislocation, and hearts were quickly excised and submerged in liquid nitrogen. Hearts were then cryopulverized, suspended in homogenization buffer (0.9% NaCl, 10 mM Tris-HCl, 20 mM NaF, 2 μM leupeptin, 100 μM phenylmethylsulphonyl fluoride, 500 μM benzamidine and 100 nM aprotinin with pH 6.8), homogenized using a Teflon pestle, and centrifuged at 1,000 x g for 8 minutes at 4 °C. Supernatants were aliquoted and stored at −80 °C until used. Bradford assays (Bio-Rad, 5000006) were used to measure protein concentration. For measuring protein expression, 50 μg of tissue homogenate were suspended in Laemmli buffer (Bio-Rad, 1610747) and separated by SDS-PAGE in 4–20% TGX (Bio-Rad, 3450064) or 16.5% Tris-Tricine precast gels (Bio-Rad, 3450033). Proteins were then transferred to PVDF membranes using the iBlot 2 transfer system (ThermoFisher) or overnight wet transfer. The following primary antibodies were used to probe membranes for specific ECC proteins: anti-RyR (clone 34C) (1:2,000; ThermoFisher, MA3-925), anti-SERCA2 (clone 2A7-A1) (1:1,000; ThermoFisher, MA3-919), anti-NCX1 (clone EPR12739) (1:1,000; Abcam, ab177952), anti-CaV1.2 (1:500; Alomone, ACC-003), and anti-PLB (clone A1) (1:5,000; Badrilla, A010-14). For RyR2 phosphorylation, custom-made antibodies were used for phospho-S2031 and phospho-S2808, while a commercial antibody was used for phospho-S2814 (1:1,000, Badrilla, A010-31AP). Anti-GAPDH (clone 6C5) (1:10,000; MilliporeSigma, MAB374) was used as a loading control. Secondary antibodies, goat anti-mouse-HRP (1:2,000; ThermoFisher, 31437) or goat anti-rabbit-HRP (1:1,000; ThermoFisher, 31463), were used as appropriate. Membranes were developed using Clarity ECL substrate (Bio-Rad, 1705061) or SuperSignal Chemiluminescent Substrate (ThermoFisher, 34580 or 34096). A ChemiDoc MP apparatus (Bio-Rad) was used to image membranes. Band intensity was quantified using ImageLab 6.1 software (Bio-Rad).

### Sarcomere contraction/relaxation recordings

Sarcomere length measurements were obtained using the IonOptix Myocyte Sarcomere Length acquisition system equipped with a MyoCam to determine cell dimension changes. Cardiomyocytes were freshly isolated and maintained in normal Tyrode solution supplemented with 1.8 mM CaCl₂. Cells were then added to a perfusion and stimulation chamber where 40 V, 10 ms pulses were delivered at 1 Hz via platinum electrodes. Recordings were conducted at room temperature. To assess cell contraction under adrenergic stimulation, we recorded under basal conditions for at least 1 min, after that Iso (100 nM) was perfused continuously for 8 min with continuous recording. Data for basal conditions was selected after sarcomere length was stable and for the Iso response, data was collected at least 5 min after Iso perfusion. Sarcomere length kinetic parameters were analyzed and quantified using IonWizard (IonOptix) analysis software.

### Echocardiography recordings

Transthoracic long-axis M-mode echocardiography was performed by the University of Wisconsin-Madison Small Animal Imaging and Radiotherapy Facility (SAIRF) at the University of Wisconsin-Madison. A Vevo 3100 system running Vevo Lab 5.8.2 (Visual Sonics) with a 22– 55-MHz transducer (Visual Sonics, MS550D) was used for all recordings. Mice were anesthetized with 1–1.5% isoflurane and maintained at a body temperature of 37.0 ± 0.5°C using a heated platform. Two-dimensionally guided M-mode images of the long axis of the left ventricle were recorded. Ejection fraction, fractional shortening, left ventricle and posterior wall metrics, and heart rate were all calculated through analysis of left ventricle diameter, septum wall thickness, and posterior wall thickness. At least three consecutive cardiac cycles were analyzed for all measurements.

### Electrocardiography (ECG) recordings

Mice were anesthetized with isoflurane (2%) and maintained at a body temperature of 37.0 ± 0.5°C with a heating pad. Needle electrodes were placed subcutaneously in each limb to record in Lead-I and Lead-II ECG configurations, using a PowerLab system and LabChart 8.1.21 (ADInstruments). After stabilization of body temperature and heart rate, mice were allocated to one of two treatment groups: (1) isoproterenol (2 mg/kg) was injected intraperitoneally to test the chronotropic response in vivo; (2) a cocktail containing epinephrine (4 mg/kg) and caffeine (120 mg/kg) was injected intraperitoneally to evaluate the susceptibility to stress-induced arrhythmia. At least 5 minutes of basal ECG were recorded followed by 10-15 min of monitoring post-injection.

### Patch-clamp recordings of I_CaL_

Whole-cell patch clamp was used to measure *I*_CaL_. Patch-clamp recordings were done using an Axopatch 200B amplifier paired with a Digidata 1550A digitizer controlled with Clampex 10 (Axon Instruments). For basal recordings, cells were kept in Tyrode’s solution (described above) supplemented with 30 μM tetrodotoxin (TTX, Cayman Chemical #14964) and 5 mM 4-aminopyridine (4-AP, Sigma #275875), to block *I*_Na_ and *I*_to_, respectively. Cells used for Iso recordings were kept in the same solution plus 100 nM isoproterenol. Borosilicate glass pipettes (World Precision Instruments, 1B150F-4) were pulled using a Flaming/Brown micropipette puller (Sutter Instruments, model P-1000). Pipette resistance was kept at 2-3 MΩ, and appropriate compensation (≥70%) for whole-cell capacitance and series resistance was applied. Pipettes were filled with an internal solution (110 mM CsCl, 6 mM MgCl_2_, 5 mM Na_2_ATP, 0.3 mM Na_2_GTP, 10 mM HEPES and 15 mM TEAꞏCl, with pH adjusted to 7.2 using CsOH). *I*_CaL_ was elicited using a holding potential of −40 mV and applying 300 ms pulses from −40 mV to 50 mV in 10 mV increments. All recordings were performed at room temperature. Current densities (pA/pF) were calculated by dividing the peak current by cell capacitance. Data analysis was performed using Clampfit 10.

### Immunoprecipitation and incorporation of ^32^P

SR-enriched mouse microsomes were prepared as previously reported.^17^ Ventricles from 10-15 mouse hearts were pooled together and homogenized as indicated above. The resulting homogenate was centrifuged at 8,000 x *g* for 20 min at 4 °C. The supernatant was collected and further centrifuged at 100,000 x *g* for 35 min at 4 °C. The pellet was resuspended in homogenization buffer containing 0.3 mol/L of sucrose, aliquoted and stored at −80 °C until used. Protein concentrations were determined using the Bradford method (Bio-Rad).

For RyR2 immunoprecipitations,^42^ 100 μg of protein from SR-enriched microsomes were diluted in lysis buffer (homogenization buffer containing 0.1% Triton X-100). Samples were incubated for 2 hours at room temperature with protein-G Dynabeads (ThermoFisher) conjugated with 2 μg of anti-RyR antibody (ThermoFisher, MA3-925), as per the manufacturer’s instructions. The beads were then washed twice with lysis buffer.

Phosphorylation reactions in the presence of [γ,^32^P]-ATP were carried on the RyR2-bound beads.^17^ After washing twice with reaction buffer, the beads were resuspended in phosphorylation buffer and incubated at 37 °C for 10 minutes. Laemmli buffer with β-mercaptoethanol (Bio-Rad) was added to terminate the reactions. Western blots were carried out with the supernatant as indicated above. ^32^P incorporation was measured in PVDF membranes used for RyR2 western blots. Membranes were dried, exposed overnight on a phospho-screen (Molecular Dynamics) and imaged using a Typhoon Phosphorimager (GE Healthcare). Band intensity was quantified with the ImageLab software (Bio-Rad) and normalized to total RyR2 for each sample quantified from the same membrane.

PKA reaction buffer contained 2 mM MgCl_2_, 60 mM KCl, 30 mM HEPES, 1 mM EGTA-Na, 20 mM NaF, 2 µM leupeptin, 100 µM phenylmethylsulphonyl fluoride, 500 µM benzamidine, 100 nM aprotinin, 20 mM NaF, 5 μM okadaic acid potassium, pH 7.4. PKA phosphorylation buffer was the reaction buffer supplemented with 10 U/μL of the catalytic subunit of PKA (539576, Millipore), 1 mM K_2_ATP and 0.3 μM [γ,^32^P]-ATP (BLU502A, PerkinElmer). CaMKII reaction buffer contained 2 mM MgCl_2_, 60 mM KCl, 30 mM HEPES, 20 μM CaCl_2_, 20 mM NaF, 2 µM leupeptin, 100 µM phenylmethylsulphonyl fluoride, 500 µM benzamidine, 100 nM aprotinin, 20 mM NaF, 5 μM okadaic acid potassium, pH 7.4. CaMKII phosphorylation buffer was the reaction buffer supplemented with 1 μL of recombinant CaMKIIδ (PV3373, ThermoFisher), 3 μM CaM (208694, Sigma), 1 mM K_2_ATP and 0.3 μM [γ,^32^P]-ATP (as above). Control reactions were carried out in PKA reaction buffer supplemented with 1 mM K_2_ATP and 0.3 μM [γ,^32^P]-ATP (as above) but without the addition of enzymes.

### Statistical analysis and reproducibility

All experiments performed at institutional cores were single blinded. Most datasets were collected over multiple days, and results were consistent within groups. All numerical results, bar graphs, time-courses, and curves are presented as mean ± SEM. Data management was done using Excel 365 (Microsoft) and Origin 2025b (OriginLab). Statistical significance was determined at p < 0.05 using the tests and sample sizes indicated in each figure legend. Comparisons of protein expression solely between genotypes were done using two-sided t-test (parametric) or Mann–Whitney rank-sum test (non-parametric) in SigmaPlot 15.0 (Systat Software). Comparisons of RyR2 phosphorylation between genotypes and treatments were done using two-way ANOVA also with SigmaPlot 15.0. All analyses for cellular and whole-animal experiments used a linear mixed-effects model (LMM) in SPSS 31.0 (IBM Corp.) to account for the hierarchical structure of data obtained in single cardiomyocytes and/or data involving repeated measurements within the same cell/animal. For CaT and sarcomere contraction/relaxation analysis, treatment and genotype were included as fixed effects, and individual mice were included as a random effect to account for hierarchical correlation. For *I*_CaL_ analysis with two data hierarchies (repeated measures at different voltages in each cell and multiple cardiomyocytes from the same mouse), treatment, genotype and voltage were included as fixed effects, individual mice and cells were included as a random effect, and voltage was included as a repeated measure. For echocardiography and electrocardiography analysis with multiple measurements within the same mouse, genotype and treatment were included as fixed effects, individual mice as a random effect, and treatment as a repeated measure. The Bonferroni adjustment/test was used for all LMM analyses.

## Acknowledgements

This work was supported by the National Institutes of Health (R01HL161070 and R01HL167195 to FJA) and start-up funds from the UW SMPH Department of Medicine (FJA). AMG was supported by the Cardiovascular Research Center Summer Undergraduate Fellowship.

We thank Ms. Holly C. Dooge, who provided technical assistance in the execution of experiments involving [γ,^32^P]-ATP, and Dr. Angela Greenman (University of Wisconsin-Madison) for providing feedback on the manuscript. We also thank Dr. Steven Marx (Columbia University) for kindly providing the 4SA-Rad mouse model.

## Author contributions

FJA conceived the project. FJA, AMG and EBRP conceptualized most experiments. FJA and AMG wrote the manuscript. AMG collected and analyzed calcium imaging data in isolated cardiomyocytes and western blot data in heart tissues and prepared the figures. EBRP collected and analyzed cell shortening and echocardiography data. DPB collected and analyzed *I_CaL_* data. FJA performed the [γ,^32^P]-ATP experiments. FJA, SLS, AMG and EBRP collected and analyzedECG data. All authors discussed the results and approved the manuscript.

## Extended Data

**Extended Data Fig. 1.**
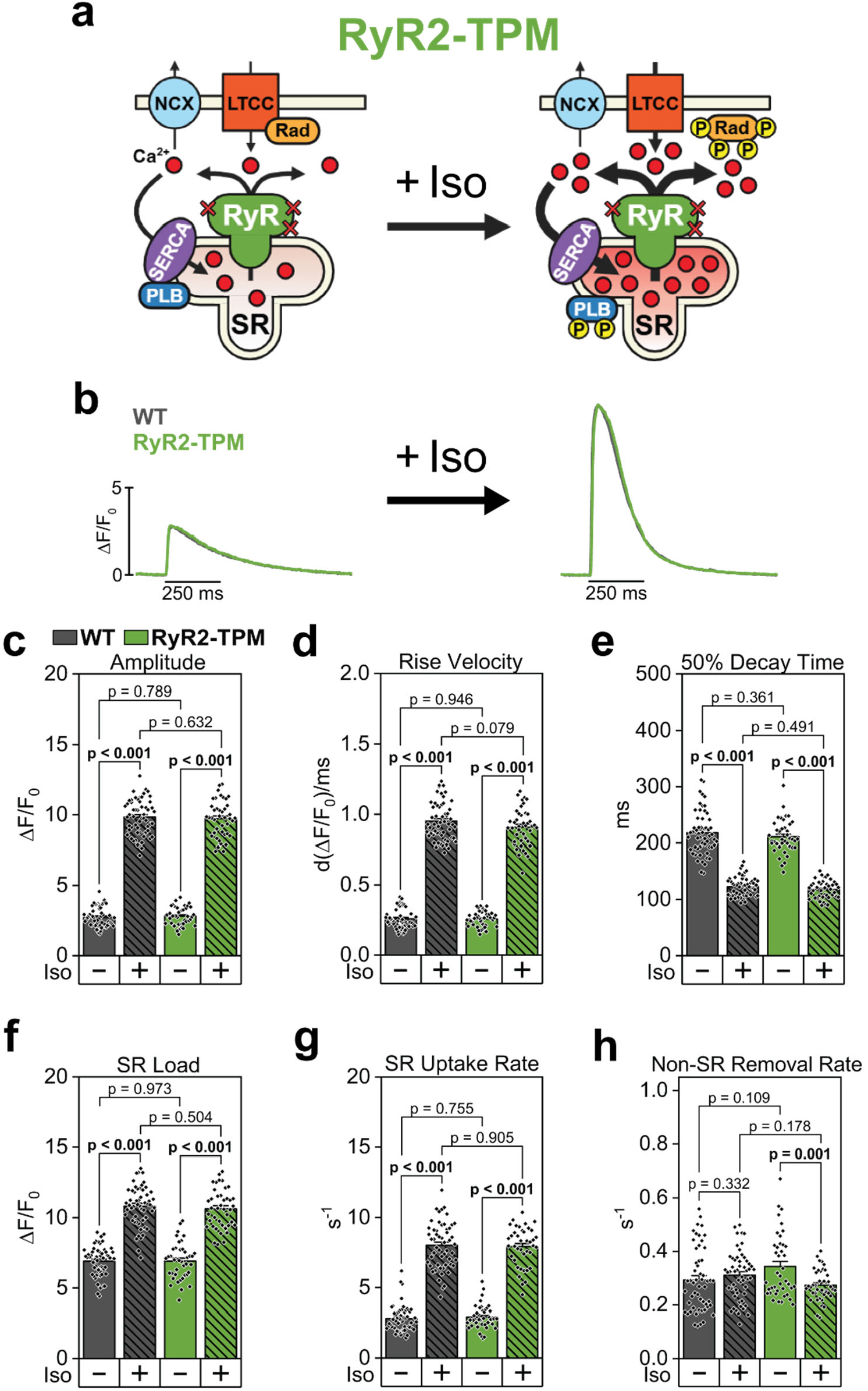
CaT properties in RyR2-TPM cardiomyocytes are indistinguishable from WT controls. **a.** Diagram of Ca^2+^ handling response to β-AR activation by Iso in RyR2-TPM cardiomyocytes. Canonical phosphorylation sites on RyR2 (S2031, S2808, and S2814) are ablated, although there is no resulting change in CaT properties. **b.** Representative confocal CaT fluorescence profiles obtained in wild-type (WT) and RyR2-TPM ventricular cardiomyocytes at 1 Hz pacing after 120 s under basal conditions or in the presence of 100 nM Iso. Data are presented as corrected fluorescence (ΔF/F_0_). **c-h.** Quantification of CaT properties at 1 Hz pacing, after 120 s in basal conditions or under 100 nM Iso perfusion, including CaT amplitude (**c**), maximum rise velocity (**d**), 50% decay time (**e**), measured as the time from the CaT peak to 50% CaT decay, SR load (**f**), measured as the peak of a caffeine-induced CaT at the end of the CaT recording protocol, SR Ca^2+^ uptake rate (**g**), and non-SR Ca^2+^ removal rate (**h**). A single exponential fitting of the decay phase of the caffeine-induced transient was used to determine non-SR Ca^2+^ removal, while the SR Ca^2+^ uptake rate was calculated as the constant from exponential fitting of paced CaT decay minus the non-SR removal constant. N = 6 mice per genotype for all panels. n = 56 WT and 42 RyR2-TPM [Basal and Iso] myocytes for panels c-e. n = 55 WT [Basal], 54 WT [Iso], and 41 RyR2-TPM [Basal and Iso] myocytes for panel f. n = 54 WT [Basal and Iso], 41 RyR2-TPM [Basal], and 40 RyR2-TPM [Iso] myocytes for panels g-h. Linear mixed-effects model (Bonferroni test) used for all comparisons.

**Extended Data Fig. 2.**
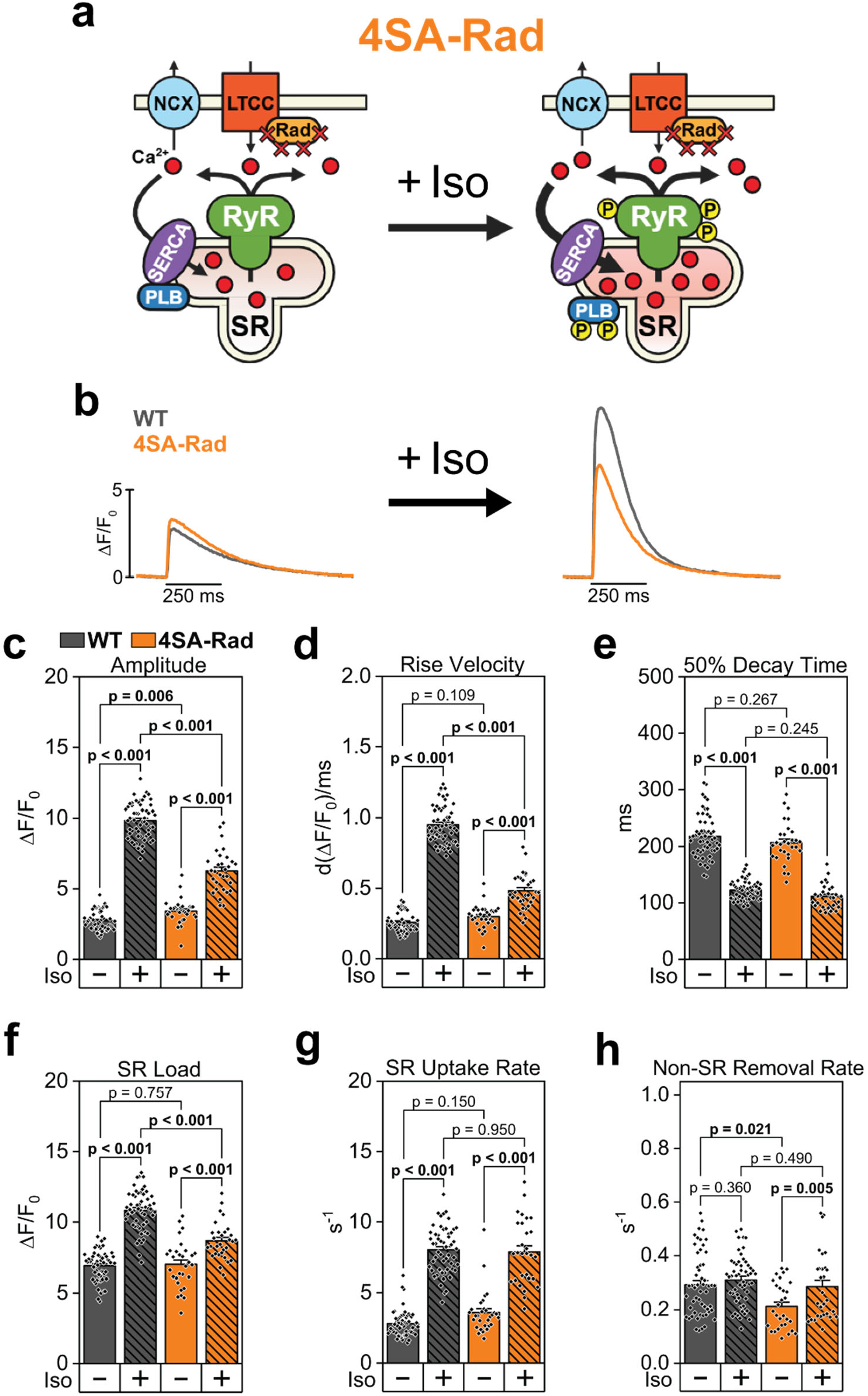
CaT properties are blunted in response to Iso in 4SA-Rad cardiomyocytes. **a.** Diagram of Ca^2+^ handling response to β-AR activation by Iso in 4SA-Rad cardiomyocytes. Phosphorylation sites on Rad (S25, S38, S272, and S300) are ablated, preventing adrenergic regulation of *I*_CaL_ through LTCC and blunting the Iso response of CaT properties. **b.** Representative confocal CaT fluorescence profiles obtained in wild-type (WT) and 4SA-Rad ventricular cardiomyocytes at 1 Hz pacing after 120 s under basal conditions or in the presence of 100 nM Iso. Data are presented as corrected fluorescence (ΔF/F_0_). **c-h.** Quantification of CaT properties at 1 Hz pacing, after 120 s in basal conditions or under 100 nM Iso perfusion, including CaT amplitude (**c**), maximum rise velocity (**d**), 50% decay time (**e**), measured as the time from the CaT peak to 50% CaT decay, SR load (**f**), measured as the peak of a caffeine-induced CaT at the end of the CaT recording protocol, SR Ca^2+^ uptake rate (**g**), and non-SR Ca^2+^ removal rate (**h**). A single exponential fitting of the decay phase of the caffeine-induced transient was used to determine non-SR Ca^2+^ removal, while the SR Ca^2+^ uptake rate was calculated as the constant from exponential fitting of paced CaT decay minus the non-SR removal constant. N = 6 mice per genotype for all panels. n = 56 WT and 30 4SA-Rad [Basal and Iso] myocytes for panels c-e. n = 55 WT [Basal], 54 WT [Iso], and 30 4SA-Rad [Basal and Iso] myocytes for panel f. n = 54 WT [Basal and Iso], 30 4SA-Rad [Basal], and 29 4SA-Rad [Iso] myocytes for panels g-h. Linear mixed-effects model (Bonferroni test) used for all comparisons.

**Extended Data Fig. 3.**
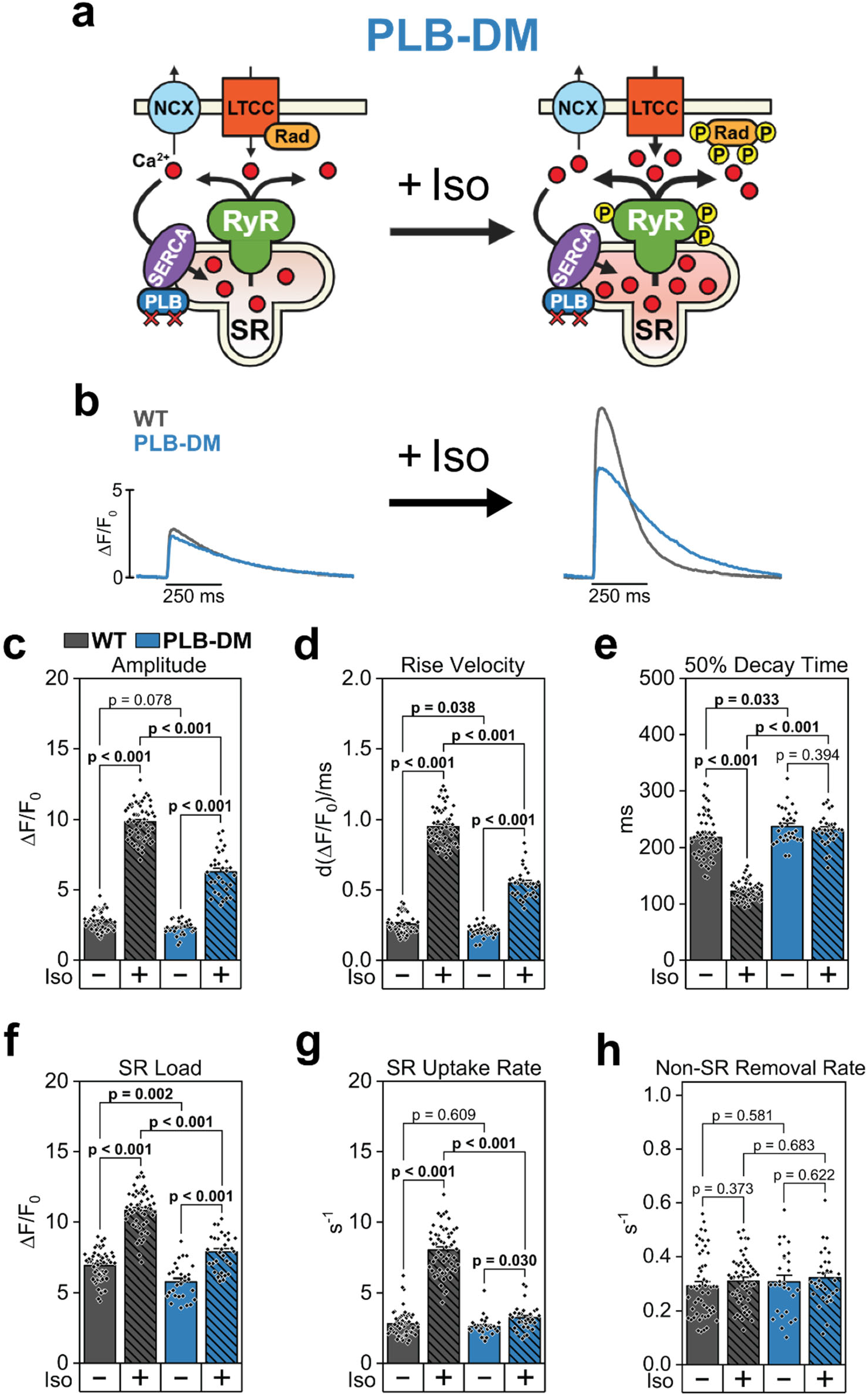
CaT properties are blunted in response to Iso in PLB-DM cardiomyocytes. **a.** Diagram of Ca^2+^ handling response to β-AR activation by Iso in PLB-DM cardiomyocytes. Phosphorylation sites on PLB (S16 and T17) are ablated, preventing direct adrenergic regulation of SR Ca^2+^ uptake by SERCA2 and blunting the Iso response of CaT properties. **b.** Representative confocal CaT fluorescence profiles obtained in wild-type (WT) and PLB-DM ventricular cardiomyocytes at 1 Hz pacing after 120 s under basal conditions or in the presence of 100 nM Iso. Data are presented as corrected fluorescence (ΔF/F_0_). **c-h.** Quantification of CaT properties at 1 Hz pacing, after 120 s in basal conditions or under 100 nM Iso perfusion, including CaT amplitude (**c**), maximum rise velocity (**d**), 50% decay time (**e**), measured as the time from the CaT peak to 50% CaT decay, SR load (**f**), measured as the peak of a caffeine-induced CaT at the end of the CaT recording protocol, SR Ca^2+^ uptake rate (**g**), and non-SR Ca^2+^ removal rate (**h**). A single exponential fitting of the decay phase of the caffeine-induced transient was used to determine non-SR Ca^2+^ removal, while the SR Ca^2+^ uptake rate was calculated as the constant from exponential fitting of paced CaT decay minus the non-SR removal constant. N = 6 mice per genotype for all panels. n = 56 WT and 30 PLB-DM [Basal and Iso] myocytes for panels c-e. n = 55 WT [Basal], 54 WT [Iso], 28 PLB-DM [Basal], and 30 PLB-DM [Iso] myocytes for panel f. n = 54 WT [Basal and Iso], 28 PLB-DM [Basal], and 30 PLB-DM [Iso] myocytes for panels g-h. Linear mixed-effects model (Bonferroni test) used for all comparisons.

**Extended Data Fig. 4.**
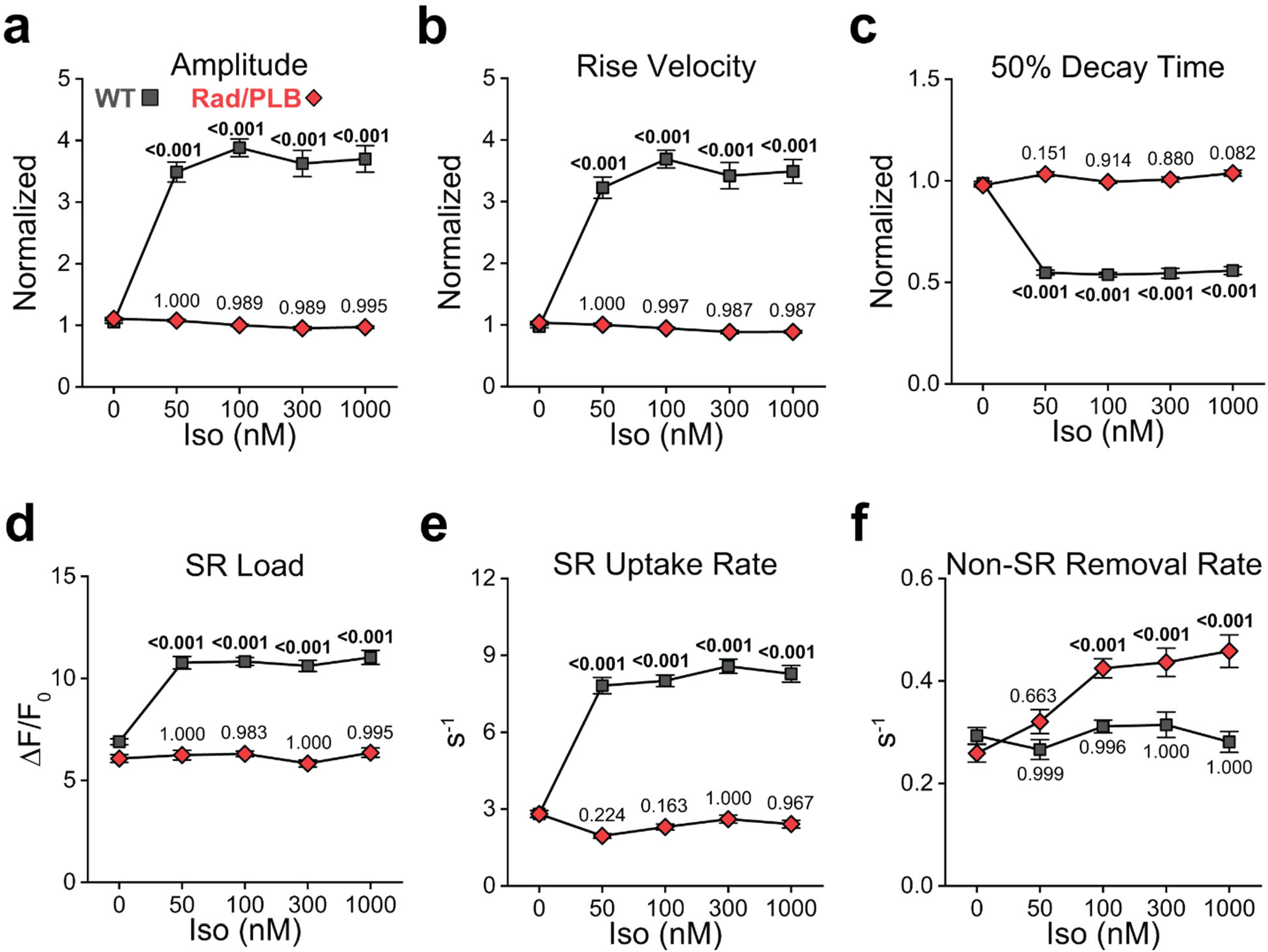
CaT properties in Rad/PLB cardiomyocytes except for non-SR Ca^2+^ uptake are unresponsive to Iso at varying concentrations. **a-f.** Quantification of CaT properties at 1 Hz pacing, after 120 s in basal conditions or under 100 nM Iso perfusion, including CaT amplitude (**a**) normalized to basal value for each cell, maximum rise velocity (**b**) normalized to basal value for each cell, 50% decay time (**c**) normalized to basal value for each cell, SR load (**d**), measured as the peak of a caffeine-induced CaT at the end of the CaT recording protocol, SR Ca^2+^ uptake rate (**e**), and non-SR Ca^2+^ removal rate (**f**). A single exponential fitting of the decay phase of the caffeine-induced transient was used to determine non-SR Ca^2+^ removal, while the SR Ca^2+^ uptake rate was calculated as the constant from exponential fitting of paced CaT decay minus the non-SR removal constant. N = 6 mice per genotype for all panels. n = 56 WT [0 and 100 nM Iso], 24 WT [50, 300, and 1000 nM Iso], 56 Rad/PLB [0 and 100 nM Iso], 23 Rad/PLB [50 nM Iso], and 24 Rad/PLB [300 and 1000 nM Iso] myocytes for panels a-c. n = 55 WT [0 nM Iso], 23 WT [50, 300, and 1000 nM Iso], 54 WT [100 nM Iso], 54 Rad/PLB [0 nM Iso], 23 Rad/PLB [50 and 300 nM Iso], 55 Rad/PLB [100 nM Iso], and 22 Rad/PLB [1000 nM Iso] myocytes for panel d. n = 54 WT [0 and 100 nM Iso], 23 WT [50, 300, and 1000 nM Iso], 54 Rad/PLB [0 and 100 nM Iso], 23 Rad/PLB [50 and 300 nM Iso], and 22 Rad/PLB [1000 nM Iso] myocytes for panels e-f. Linear mixed-effects model (Bonferroni test) used for all comparisons. P-values for comparisons within genotype to basal (0 nM Iso) shown directly above or below data point.

**Extended Data Fig. 5.**
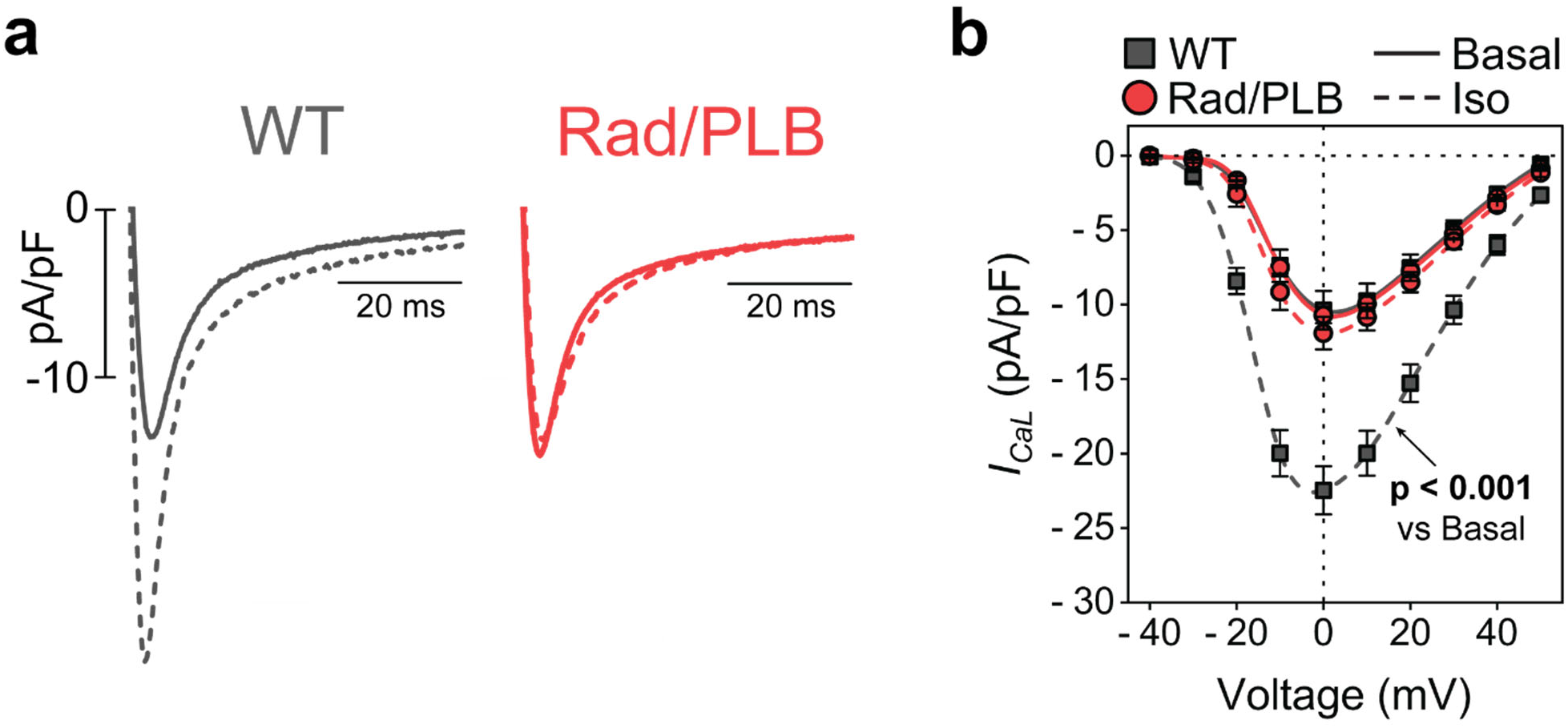
*I*_CaL_ in Rad/PLB cardiomyocytes does not respond to β-adrenergic stimulation with Iso. **a.** Representative voltage clamp recordings of *I*_CaL_ at 0 mV with or without 100 nM Iso. **b.** Current-voltage (I-V) curves of *I*_CaL_ density with Boltzmann fitting of experimental data shown as solid (basal) and dashed (Iso) lines. N = 5 WT and 4 Rad/PLB mice. n = 13 WT [Basal], 14 WT [Iso], 17 Rad/PLB [Basal], and 15 Rad/PLB [Iso] myocytes. Linear mixed-effects model (Bonferroni test) with repeated measures. P < 0.001 between WT Basal and WT Iso curves.

**Extended Data Fig. 6.**
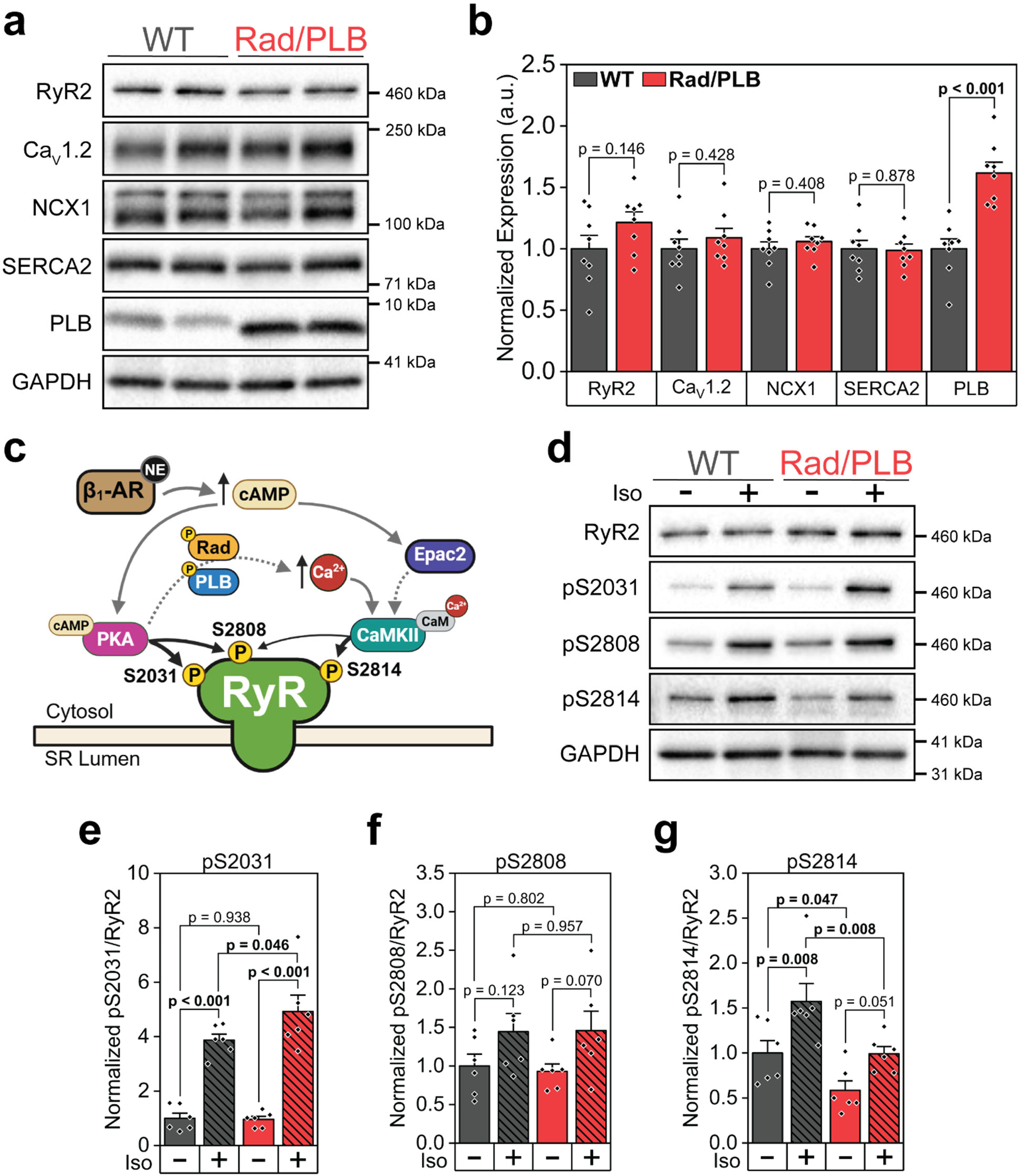
Rad/PLB hearts show little alteration of Ca^2+^-handling protein expression while RyR2 remains phosphorylated. **a.** Representative western blot images for major excitation-contraction coupling (ECC) proteins involved in control of Ca^2+^ dynamics: ryanodine receptor 2 (RyR2), the L-type Ca^2+^ channel (LTCC) isoform Ca_V_1.2, Na^+^/ Ca^2+^ exchanger 1 (NCX1), sarco-(endo)-plasmic reticulum Ca^2+^ ATPase 2 (SERCA2), and phospholamban (PLB). **b.** Quantification of Ca^2+^-handling protein expression in WT and Rad/PLB hearts, normalized to GAPDH. Results are shown relative to WT expression for each protein. N = 8 hearts per genotype. T-test was used for all comparisons. **c.** Diagram of RyR2 phosphorylation downstream of β-AR activation. Protein kinase A (PKA) is activated by cyclic AMP (cAMP) and phosphorylates serines 2031 and 2808 on RyR2. Ca^2+^/calmodulin-dependent kinase II (CaMKII) is activated by elevated cytosolic Ca^2+^ and through the exchange protein activated by cAMP 2 (Epac2) pathway. CaMKII mainly phosphorylates serine 2814 and partially contributes to phosphorylation of serine 2808. **d**. Representative western blot images of RyR2 and RyR2 phosphorylation at canonical sites (S2031, S2808, and S2814) in hearts collected from mice at rest or 10 min after intraperitoneal (i.p.) injection of Iso (2 mg/kg). **e-g.** Quantification of RyR2 phosphorylation at residues S2031 (**e**), S2808 (**f**), and S2814 (**g**), normalized first to GAPDH and then to RyR2 density. Results are shown relative to WT. N = 6 hearts per genotype [Basal and Iso]. Two-way ANOVA was used for all comparisons.

**Extended Data Fig. 7.**
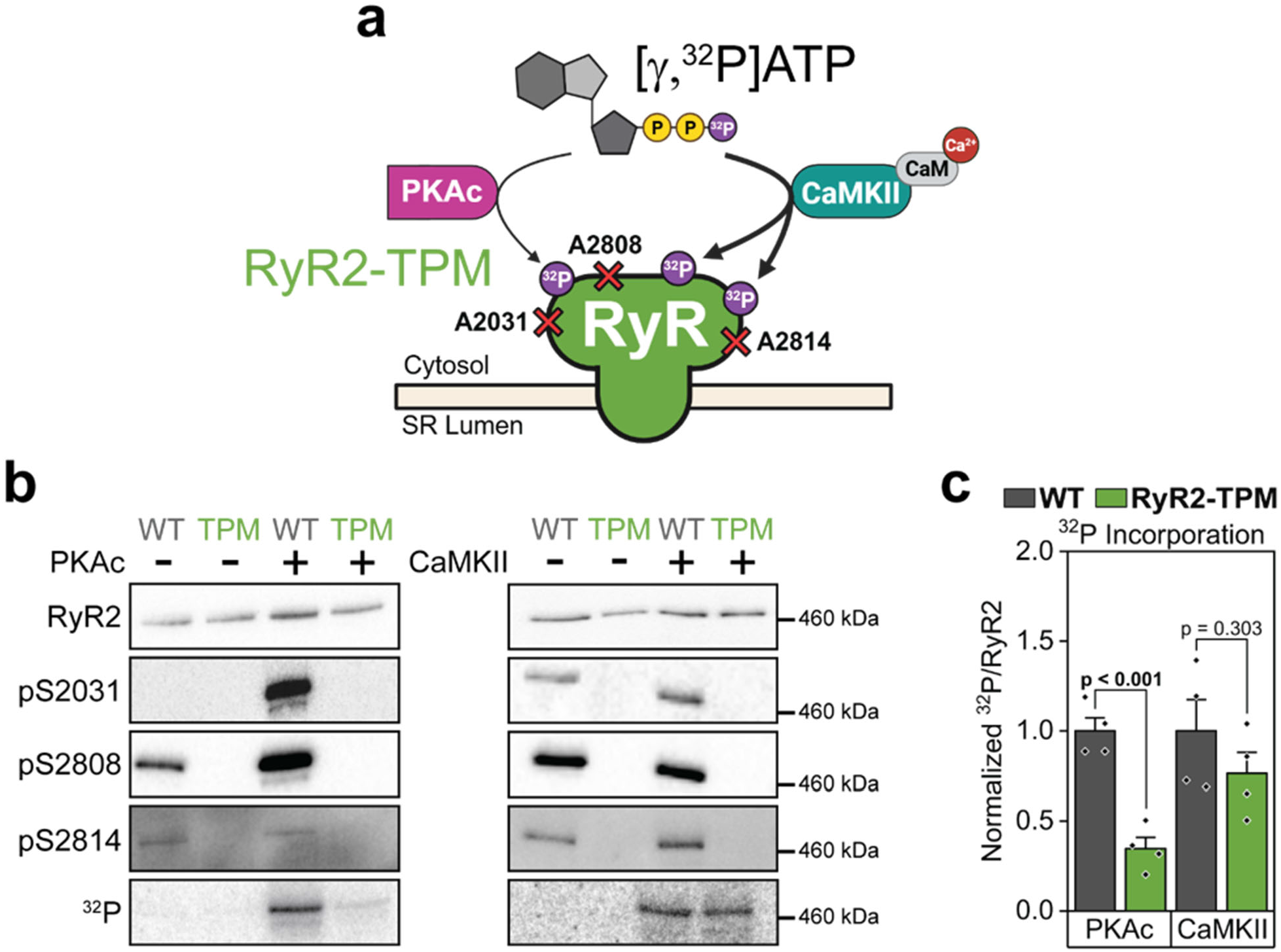
PKA and CaMKII can incorporate ^32^P into RyR2-TPM channels in vitro. **a.** Diagram of in vitro phosphorylation of RyR2-TPM channels in the presence of [γ,^32^P]-ATP. Incorporation of ^32^P into RyR2-TPM channels indicates phosphorylation of non-canonical sites. **b**. Representative images of western blots of total RyR2 and phospho-sites, and ^32^P incorporation of RyR2 immunoprecipitated from SR-enriched cardiac microsomes and phosphorylated in vitro with the catalytic subunit of PKA (PKAc) or CaMKII in the presence of [γ,^32^P]-ATP. **c**. Quantification of ^32^P incorporation into WT and RyR2-TPM channels. PKAc incorporated significantly less ^32^P into RyR2-TPM than into WT channels, while there was no significant difference in ^32^P incorporation by CaMKII (N = 4 per group. T-test used for both comparisons).

**Extended Data Table 1.** Summary of structural echocardiographic parameters in WT and Rad/PLB mice. Structural parameters measured in WT and Rad/PLB mice using long-axis M-mode echocardiography. PWd: posterior wall thickness in diastole; PWs: posterior wall thickness in systole; LVDd: left ventricle diameter in diastole; LVDs: left ventricle diameter in systole; LVVd: left ventricle volume in diastole; LVVs: left ventricle volume in systole; LV Mass Corr: corrected left ventricle mass. T-test, except parameters indicated by ^: Rank Sum Test. Significant differences (p < 0.05) highlighted in yellow. Strong tendencies (p < 0.10) highlighted in green.

|  | WT | Rad/PLB | p-value |
| --- | --- | --- | --- |
| N | 16 (8M, 8F) | 16 (8M, 8F) | - |
| PWd (mm) | 0.81 ± 0.03 | 0.82 ± 0.07 | 0.175^ |
| PWs (mm) | 1.14 ± 0.04 | 1.08 ± 0.07 | 0.142^ |
| LVDd (mm) | 3.67 ± 0.06 | 3.93 ± 0.12 | 0.050^ |
| LVDs (mm) | 2.62 ± 0.08 | 2.95 ± 0.15 | 0.078 |
| LVVd (μL) | 58.02 ± 2.43 | 68.78 ± 2.43 | 0.052 |
| LVVs (μL) | 25.66 ± 1.90 | 35.95 ± 4.08 | 0.038 |
| LV Mass Corr (mg) | 85.99 ± 3.78 | 85.11 ± 3.84 | 0.875 |

**Extended Data Table 2.** Summary of echocardiographic parameters in WT and Rad/PLB mice in response to Iso. Functional parameters measured in WT and Rad/PLB mice in basal conditions and 2 min. after i.p. injection of Iso (2 mg/kg) using long-axis M-mode echocardiography. HR: heart rate; FS: fractional shortening; EF: ejection fraction; SV: stroke volume; CO: cardiac output. Linear mixed-effects model (Bonferroni test) with repeated measures used for all comparisons. P-value within genotype represents level of significance between basal and Iso measurements within either WT or Rad/PLB mice. P-value within treatment represents level of significance between WT and Rad/PLB measurements within either basal or Iso conditions. Significant differences (p < 0.05) highlighted in yellow.

|  | WT |  | Rad/PLB |  | p-value<br>Basal vs. Iso |  | p-value<br>WT vs. Rad/PLB |  |
| --- | --- | --- | --- | --- | --- | --- | --- | --- |
| N | 16 (8M, 8F) |  | 16 (8M, 8F) |  | - |  | - |  |
|  | Basal | Iso | Basal | Iso | WT | Rad/<br>PLB | Basal | Iso |
| HR (bpm) | 498.34 ± 12.60 | 594.24 ± 7.04 | 407.81 ± 11.39 | 505.85 ± 8.35 | < 0.001 | < 0.001 | < 0.001 | < 0.001 |
| EF (%) | 56.18 ± 2.06 | 92.92 ± 1.23 | 50.57 ± 3.12 | 42.26 ± 4.39 | < 0.001 | 0.001 | 0.200 | < 0.001 |
| FS (%) | 29.02 ± 1.35 | 66.49 ± 2.36 | 25.84 ± 1.96 | 21.27 ± 2.60 | < 0.001 | 0.043 | 0.310 | < 0.001 |
| SV (μL) | 32.36 ± 1.71 | 31.52 ± 1.30 | 32.83 ± 1.42 | 26.14 ± 1.77 | 0.598 | < 0.001 | 0.837 | 0.023 |
| CO (mL/min) | 16.26 ± 1.01 | 18.79 ± 0.93 | 13.28 ± 0.52 | 13.18 ± 0.90 | 0.004 | 0.910 | 0.026 | < 0.001 |

